# Conditional Upregulation of KCC2 selectively enhances neuronal inhibition during seizures

**DOI:** 10.1101/253831

**Authors:** CS Goulton, M Watanabe, DL Cheung, KW Wang, T Oba, A Khoshaba, D Lai, H Inada, K Eto, K Nakamura, JM Power, TM Lewis, GD Housley, H Wake, J Nabekura, AJ Moorhouse

## Abstract

Efficacious neuronal inhibition is sustained by the neuronal K^+^Cl^-^ co-transporter KCC2, and loss of KCC2 function through injury or mutation is associated with altered GABAergic signalling and neuronal seizures. Here we report a transgenic mouse with conditional KCC2 overexpression that results in increased membrane transport function. Increased KCC2 has little impact on behavioural and *in vitro* assays of neuronal excitability and GABA_A_ receptor responses under resting conditions. In contrast, increased KCC2 imparts resistance to seizure-like neuronal activity in hippocampal slices and prevents the progression of mice into behavioural status epilepticus following multiple kainic acid doses. Our results demonstrate a transgenic mouse to facilitate investigations into the role of KCC2 in brain function, and provide a proof of principle that targeting KCC2 may be an effective way to selectively enhance neuronal inhibition to mitigate against diseases that involve an imbalance between excitation and inhibition.

## Introduction

Brain function depends on the appropriate balance of neuronal excitation and neuronal inhibition across neural networks. Efficacious hyperpolarizing neuronal inhibition at γ-amino-butyric acid (GABA_A_) receptors relies on Cl^-^ influx secondary to the capacity of the K-Cl-cotransporter (KCC2) to maintain a low resting intracellular Cl^-^ concentration ([Cl^-^]i) (Payne *et al*., 1996; Kaila *et al*., 2014). KCC2 is, however, dynamic in its expression levels and function, and therefore in its contribution to [Cl^-^]i homeostasis and neuronal inhibition. In developing neurons functional KCC2 is minimal and activation of GABA receptors can elicit Cl^-^ efflux and a resultant depolarisation (Rivera *et al*., 1999; Ben-Ari *et al*., 2007; Inada *et al*., 2011). A conversion towards this immature, excitatory GABA signalling phenotype is seen in adult neurons when KCC2 function decreases following brain injuries or traumas (Kahle *et al*., 2008; Kaila *et al*., 2014). A particularly strong association is evident in both rodent models and in humans, between reduced KCC2 function and impaired inhibitory control in the pathophysiology of epilepsy and seizures. The excessive neuronal activity associated with seizures causes a loss of KCC2 transport function in rodent models (Rivera *et al*., 2004; Fiumelli *et al*., 2005; Pathak *et al*., 2007; Wake *et al*., 2007; Silayeva *et al*., 2015) and consistently, some neurons within the epileptic foci of excised temporal lobe tissues from drug refractory epilepsy patients show reduced KCC2 function and less efficacious GABA_A_ receptor-mediated inhibition (Cohen *et al*., 2002; Huberfeld *et al*., 2007). The possibility that this reduced KCC2 may actually be causative for seizures is suggested by the high seizure susceptibility seen in transgenic mice with reduced KCC2 expression (Woo *et al*., 2002; Tornberg *et al*., 2005), and furthermore by the recent discovery of loss of function mutations in KCC2 in families with different forms of hereditary epilepsies (Kahle *et al*., 2014; Puskarjov *et al*., 2014; Stodberg *et al*., 2015; Saitsu *et al*., 2016). KCC2 also has transport-independent actions to stabilize dendritic spines (Li *et al*., 2007) and these epilepsy-associated mutations may both reduce Cl^-^ extrusion capacity and/or impair spine stability (Puskarjov *et al*., 2014; Saitsu *et al*., 2016). Hence, deficits in KCC2 may confer an increased susceptibility to seizure initiation, and seizure activity may be further sustained through activity induced KCC2 downregulation.

A simple conclusion from the inverse correlation between KCC2 function and seizures is that enhancing KCC2 function could potentially mitigate against seizure induction in susceptible individuals, and circumvent a vicious cycle of seizure induced loss of efficacious neuronal inhibition. Indeed, the signalling pathways which modulate KCC2 activity, and the protein itself, have been suggested as key targets for the treatment of drug refractory epilepsy (Silayeva *et al*., 2015; Kahle *et al*., 2016). KCC2 enhancers may also find therapeutic value in various brain traumas associated with altered Cl^-^ homeostasis, including axotomy, blunt trauma, ischemia and chronic pain (Wu *et al*., 2016). Such Cl^-^ extrusion enhancing drugs have been shown to restore impaired neuronal inhibition in spinal cord circuits of peripheral nerve injured mice and relieve symptoms of chronic pain and spasticity (Gagnon *et al*., 2013; Liabeuf *et al*., 2017). However, the effects of specifically and directly enhancing KCC2 expression or function in the forebrain, and how this impacts on seizures *in vivo*, has yet to be evaluated. Given the myriad of changes in signalling pathways and ion concentrations that occur in hyperactive neural circuits *in vivo*, it may be over-simplistic to assume that agents that act to increase KCC2 in cellular models can similarly reduce [Cl^-^] and enhance inhibition in the epileptic brain. Activity-dependent post translational modifications of KCC2 activity may turn off KCC2 transport (Rivera *et al*., 2004; Fiumelli *et al*., 2005; Kahle *et al*., 2013), and negate efforts to increase KCC2 expression and function. In addition, extracellular K^+^ increases during seizures and hyperexcitability may even reverse the direction of KCC2 mediated Cl^-^ transport, facilitating depolarizing GABA responses via HCO_3^-^_ and Cl^-^ efflux (Kaila *et al*., 2014). In such a scenario, the loss of KCC2 during seizures may be an adaptive response to protect against the build-up of external K+ and/or [Cl^-^]i (and water) loading. Although a detailed modelling study has suggested KCC2 downregulation contributes to exacerbating *in vivo* seizures (Buchin *et al*., 2015), in hippocampal slice models of seizures, blocking KCC2 can reduce extracellular K+ accumulation and seizure-like after-discharges (Viitanen *et al*., 2010; Hamidi & Avoli, 2015).

In short, it is very unclear as to the *in vivo* functional consequences on neuronal inhibition and seizure phenotypes in response to increased KCC2. To directly address how KCC2 modulation impacts on adult brain function and excitability, we generated a transgenic mouse where KCC2 could be overexpressed. Given that precocious overexpression of KCC2 can alter the typical patterns of neural circuit development, it was important to utilize a conditional transgenic mouse strategy in which we had temporal control of KCC2 overexpression in adult brain. Our simple hypothesis was that overexpression of KCC2 would increase neuronal Cl^-^ transport function and enhance neuronal inhibition, and we evaluated this using both *in vitro* and *in vivo* assays of neuronal excitability. Our results characterise this important mouse model, and demonstrate that enhanced KCC2 can selectively enhance GABAergic neuronal inhibition in hyperactive circuits, and can thereby mitigate against the severity of seizures in some *in vivo* models.

## Results

### Conditional overexpression of KCC2 in the transgenic mice

To investigate the impact of KCC2 overexpression on neural circuit function in the adult brain we utilized a conditional transgenic mouse strategy to allow temporal control of KCC2 expression. We employed the tetracycline inducible gene expression system (Gossen & Bujard, 1992) and generated a Tet-Off KCC2 mouse, in which KCC2 overexpression was driven by withdrawing doxycycline (Dox) from the diet (Figure 1A). The tetracycline operator construct (tetO) was inserted immediately upstream of the KCC2 translation initiation site (KCC2-tetO mice), and these KCC2-teto mice were crossed with the CAMKIIα-tTA mice (Mayford *et al*., 1996), where the tetracycline trans activator sequence (tTA) was expressed under control of the Ca-Calmodulin kinase 2α promotor. The transgenic mice could express either or both of the tTA-CaMKIIα (tTA) and KCC2-tetO (KCC2) alleles, but only in mice expressing both alleles (tTA-KCC2) was KCC2 overexpression inducible by the absence of Dox. Mice were raised on Dox-laced chow, and within days after withdrawing Dox from the diet, marked overexpression of KCCC2 mRNA was observed in the forebrain in tTA-KCC2 mice (Figure 1B). At the protein level, withdrawal of Dox for 7 days resulted in a two to three-fold increase in KCC2 in cortex, amygdala and hippocampus as compared to wild type C57Bl6 mice, or tTA-KCC2 mice maintained on Dox (Figure 1C). Increases in KCC2 protein were apparent after 2 days off Dox and appeared to continue to increase with prolonged Dox withdrawal (Supplemental Figure 1). The increase in KCC2 protein expression levels in tTA-KCC2 mice after Dox withdrawal was gradually reversed once the mice were returned to a diet containing Dox (Figure 1C).

**Figure 1.**
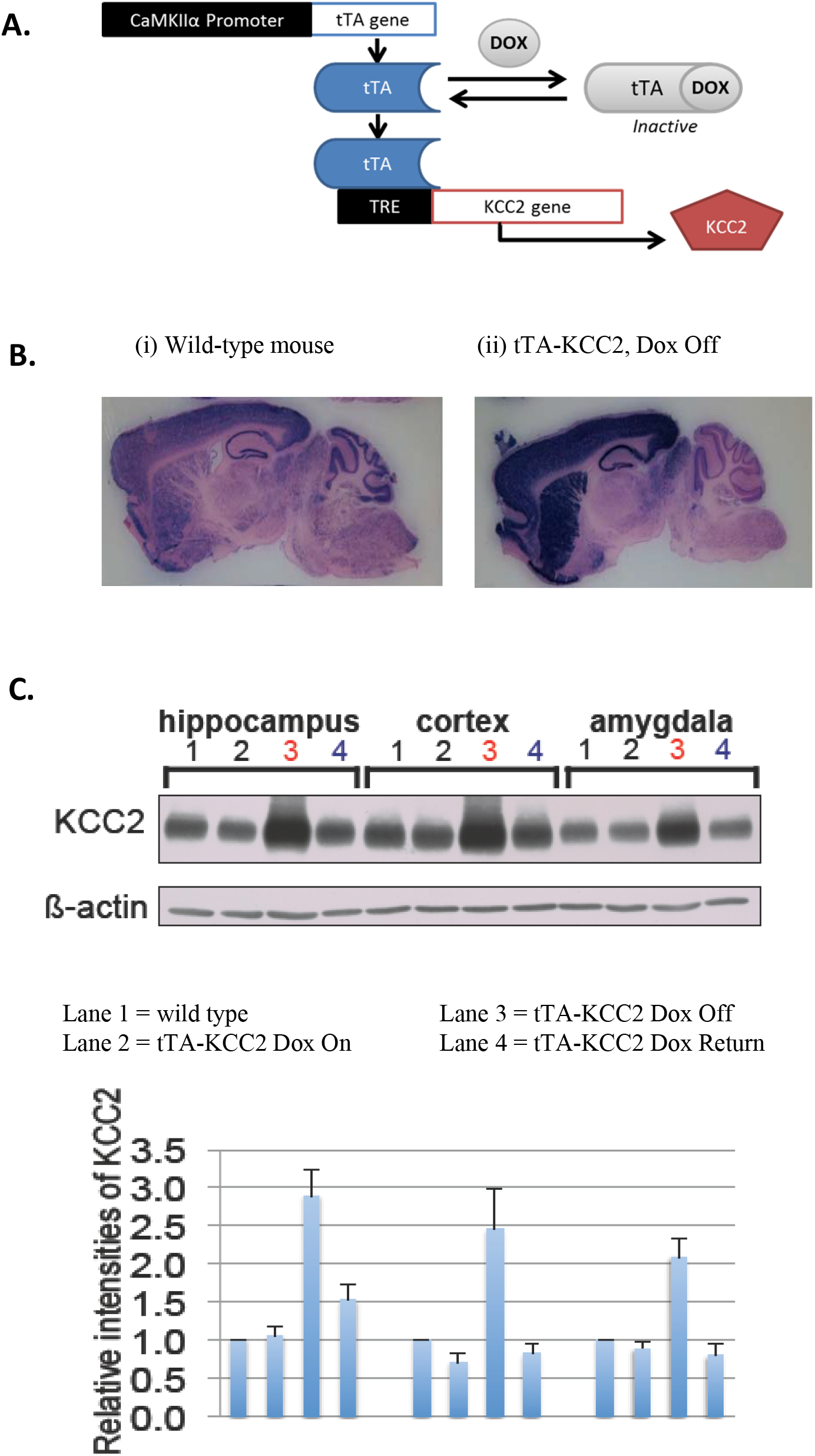
*A conditional transgenic mouse that overexpresses KCC2.* **A.** The tTA-CAMKIIa mouse was crossed with a novel transgenic mouse in which the tetO tetracycline responsive expression cassette was inserted immediately upstream of the KCC2 translation initiation site. Conditional overexpression of KCC2 in neurons expressing tTA was achieved upon withdrawal of Doxycycline (Dox) from the diet. **B.** In situ hybridization of KCC2 in sagital slices of a control, wild type (WT) C57Bl6 mouse and a transgenic mouse positive for expression of tTA-CaMKIIa and KCC2-tetO (+/+) that had Dox withdrawn from the diet for 48 hrs. **C.** Western blot for KCC2 in cortex, hippocampus and amygdala in wild type control mice (lane 1), and in tTA-KCC2 (+/+) mice continuously fed with Dox (lane 2), in which Dox was withdrawn for 7 days (lane 3) and in which Dox was withdrawn for 7 days and then reintroduced for 21 days (lane 4). Left panel shows representative blots for KCC2 and β-actin, while the right panel shows the mean data for the ratio of KCC2 to β-actin expression, normalised to that seen in the WT tissues, n = 3 different mice for each lane.

### Overexpression of KCC2 confers increased membrane KCC2 transport function

A significant proportion of KCC2 can exist in cellular localisations distinct from the membrane pools of functional oligomeric KCC2 (Watanabe *et al*., 2009). Furthermore, KCC2 function is regulated by phosphorylation (Kahle *et al*., 2013), and hence an increase in protein expression levels does not necessarily imply an increase in membrane transport capacity. To quantify KCC2 transport, we used NH_4_Cl application coupled with a fluorescence assay using the pH-sensitive membrane permeant fluorophore, BCECF-AM. Bath application of NH_4_Cl (10 mM, 3-5 minutes) to hippocampal slices from tTA-KCC2 mice that had been pre-loaded with BCECF, caused an initial alkalinisation due to passive influx of NH_3_ and a corresponding increase in BCECF fluorescence (Boron and De Weer, 1976). KCC2 can also transport NH_4_^+^ as a surrogate for K^+^, and active influx of NH_4_^+^ acidifies the internal pH resulting in decreased BCECF fluorescence (Boron & De Weer, 1976; Titz *et al*., 2006). Importantly, this assay challenges KCC2 with an imposed ionic load, rather than simply measuring resting Cl^-^ or E_GABA_, which may be a less sensitive measure of KCC2 function. All bath solutions in these experiments contained bumetanide (10 μM) to reduce any NKCC1 contribution to the fluorescent response. Examples of single neuron responses to NH_4_Cl are shown in Figure 2B. Repeated application of NH_4_Cl in the presence of furosemide (0.5 mM) reduced the acidification component of the fluorescent response, consistent with a KCC2-mediated transport (Fig. 2B). A significantly greater acidification following NH_4_Cl perfusion was seen in Dox off tTA-KCC2 mice as compared to Dox on tTA-KCC2 mice (Figure 2C), indicating that overexpression of KCC2 in Dox off mice is associated with an increase in KCC2 mediated membrane transport function.

**Figure 2.**
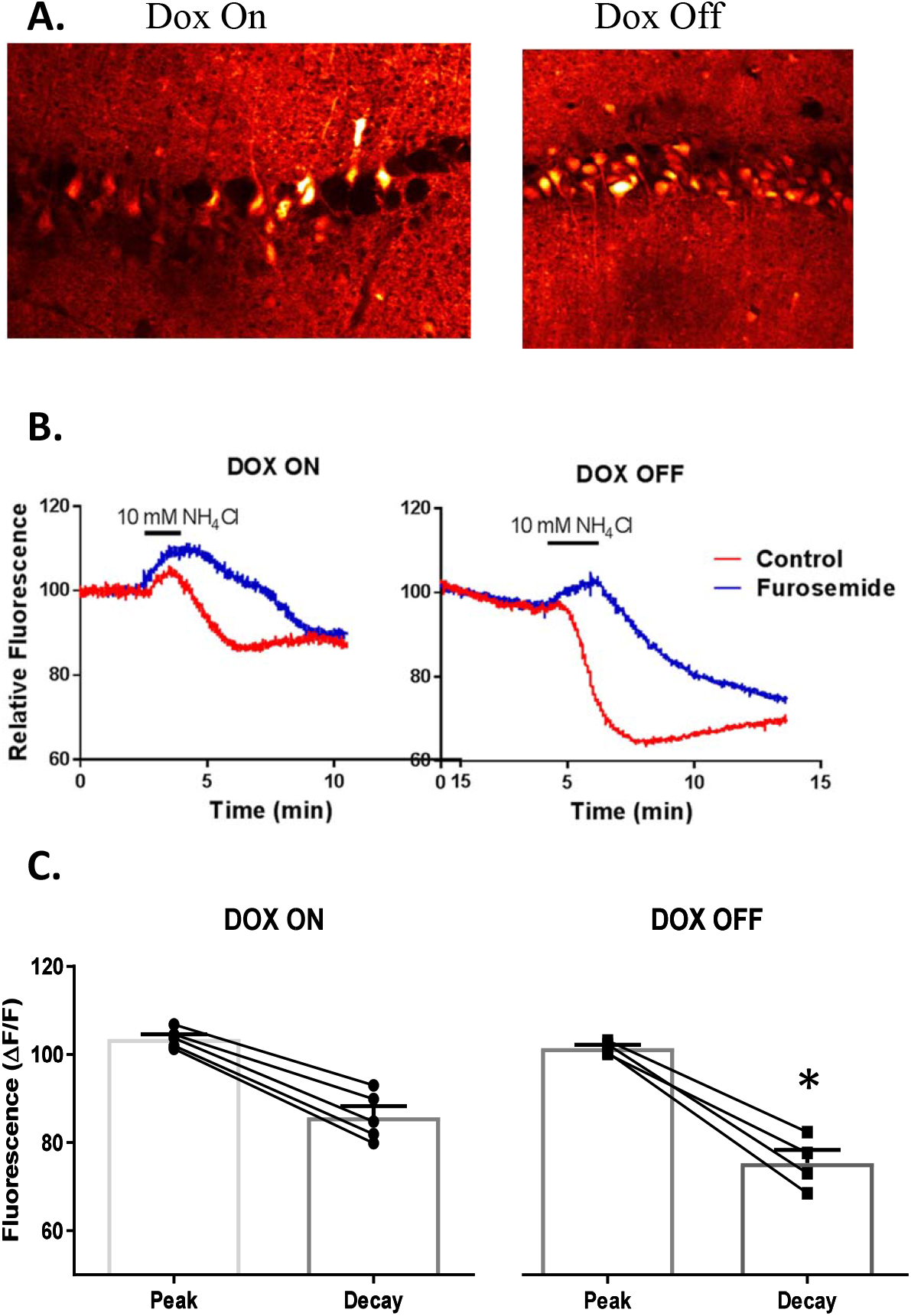
*KCC2 membrane transport function measured using BCECF fluorescence.* **A.** Example of loading and baseline fluorescence in a hippocampal slice from tTA-KCC2 mice maintained on Dox (Dox On, left) or with Dox withdrawn from the diet for 7 days (Dox Off, right). **B.** Changes in BCECF fluorescence intensity in single neurons in response to bath application of NŒUCl (10 mM) in control conditions, and then 10 minutes later in the presence of furosemide. The fluorescence has been normalised to the average of that observed in the 1^st^ 30 secs of the recording. The data come from neurons shown in **A. C.** Averaged changes in normalised fluorescence in response to NFUCl. The graphs plot the peak increase in FI and the KCC2-mediated acidification decay, measured 2 minutes after returning to control bath solution. The bar graph shows mean ± SEM, while the points and connecting lines show data from each individual experiment, * = significantly different between Dox on mice (n=5) and Dox Off mice (n=4; unpaired t-test, p = 0.03).

### Increased KCC2 does not impact on locomotion and exploratory behaviours

The open field and elevated plus maze assays are frequently used to evaluate agents which enhance neuronal inhibition, and should specifically detect sedative and/or anxiolytic effects (Kralic *et al*., 2002). As shown in Figure 3A, there was no significant difference in the total distance moved in either assay for the tTA-KCC2 Dox on vs Dox off cohorts, nor amongst any of the control or test groups. Similarly, there was no significant differences between the different mouse groups in the amount of time spent in the centre of the open field, nor the proportion of entries into the open arms of the elevated plus maze, two behaviours characteristic of anxiolytic drugs. Although withdrawal of Dox in tTA-KCC2 mice induces increased KCC2 expression and membrane transport, it doesn’t induce obvious impairments in activity levels, nor does it induce any anxiolytic effects in these assays.

**Figure 3.**
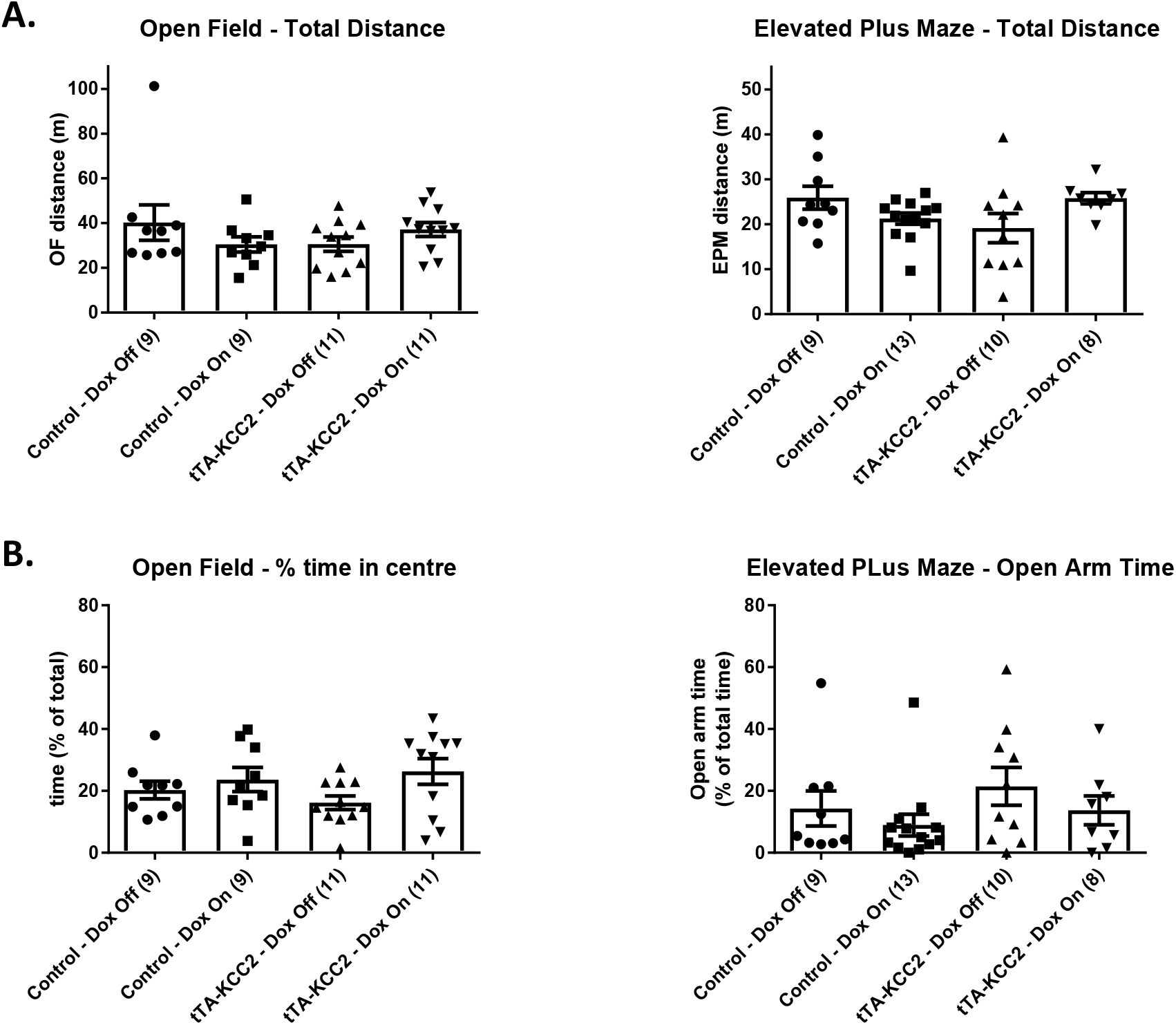
*Effects of KCC2 overexpression on locomotion and exploratory behaviours.* A. Total distanced travelled during 10 mins in an open field (left) and during 5 minutes in the elevated plus maze test (right). B. Proportion of the time spent in the centre area during 10 mins in the open field (right panel), and proportion of time spent in the open arms during 5 mins in the elevated plus maze test (right). Control refers to combined data for both Wild type (tTA-KCC2 negative) and KCC2 (tTA negative) mice, and sample size is shown in parenthesis after each group. Column graphs show mean and SEM, while individual symbols represent individual data points. There were no significant differences between mouse groups in any of these parameters (Kruskal-Wallis, and Dunn’s pot-hoc comparison tests).

### Basic properties of synaptic transmission and muscimol responses in hippocampal slices

To examine if upregulation of KCC2 alters neuronal excitability under basal or resting conditions we utilized field potential recordings in hippocampal slices, enabling the evaluation of population responses in neurons with undisturbed ionic milieu. The synaptically-evoked population response in mice with upregulated KCC2 (tTA-KCC2 mice, Dox Off) appeared the same as in three different control mice cohorts (tTA-KCC2 mice, Dox on, and KCC2 mice with or without Dox), and there was no differences amongst the groups in regards to the stimulus-response relationship, either at room temperature (22-24°C), or at an elevated bath temperature (30-32°C; Fig 4A). This suggests that neurons with increased KCC2 did not require stronger synaptic inputs to depolarize them to their action potential threshold, as may be predicted if, for example, they were more hyperpolarized at rest. Application of the GABA_A_ receptor agonist muscimol led to a concentration-dependent and reversible decrease in the amplitude of the population spike (Figure 4B). The muscimol concentration-response curves for all four experimental groups were averaged and then fit with the Hill equation to derive IC50s of 1.5 ± 1.2 (μM, mean ± SEM) for the KCC2 Dox On mice; 1.4 ± 1.2 for the KCC2 Dox Off mice; 1.4 ± 1.2 for the tTA-KCC2 Dox On mice; and 0.9 ± 1.2 for the tTA-KCC2 Dox Off, this latter group representing increased KCC2 expression. A post-hoc comparison between tTA-KCC2 Dox Off and tTA-KCC2 Dox On revealed no significant difference in IC50s (ANOVA, p=0.04; Bonferroni’s test: p=0.16). Hence upregulation of KCC2 did not cause a significant increase in the muscimol potency.

**Figure 4.**
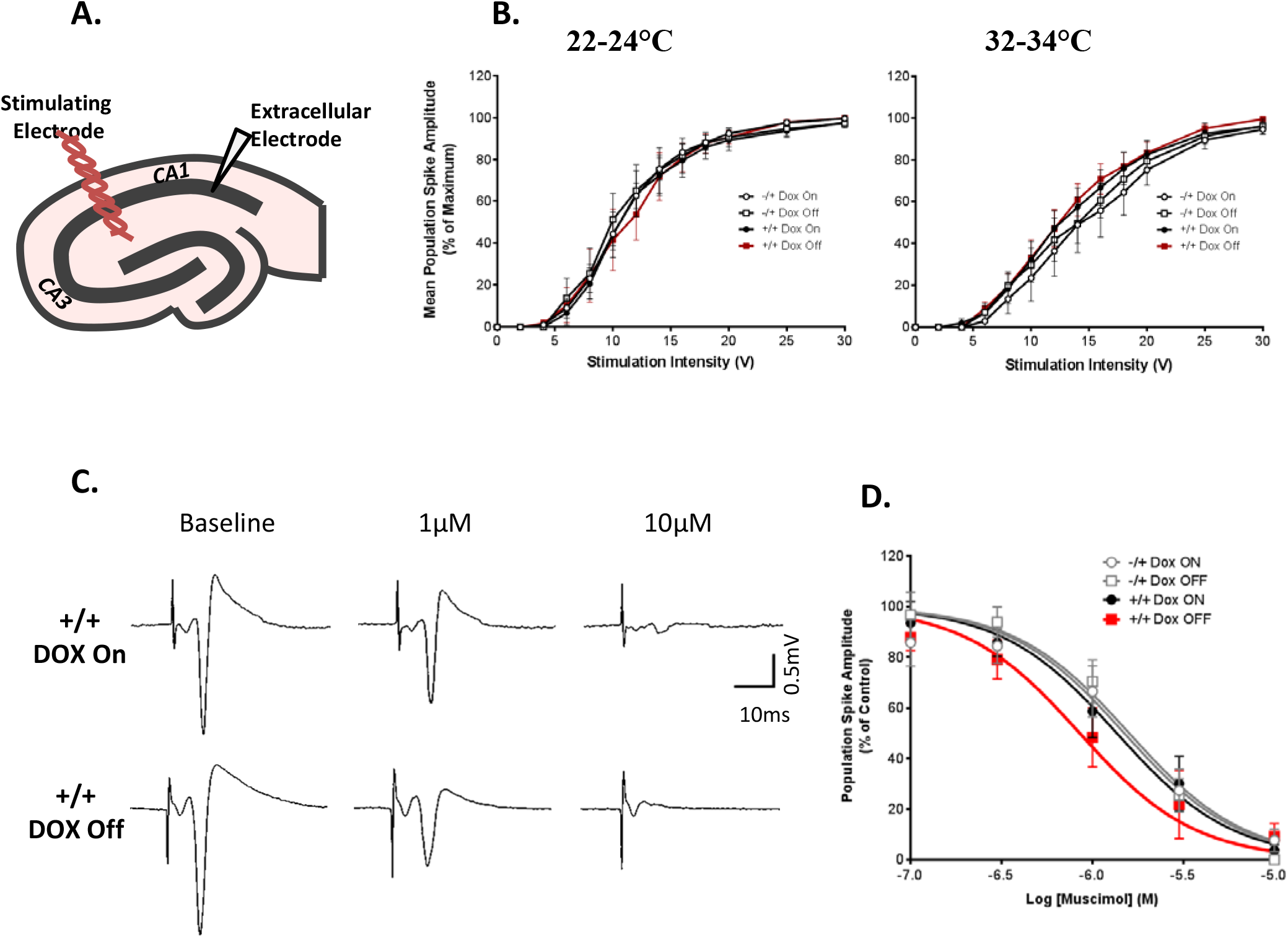
*Basal properties of field potentials and GABAA-receptor mediated inhibition in hippocampal slices from transgenic mice.* **A.** Schematic illustration of experimental setup. The population spike was recorded from the CA1 pyramidal cell layer in response to stimulation of Schaeffer collateral afferents arising from CA3 neurons. **B.** Increasing stimulus intensity resulted in an increasing population spike amplitude up until a maximum. There was no difference in this input-output relationship between any of the mouse groups at either room temperature (left panel) or at elevated temperature (right panel). Population spike amplitudes were normalised to those obtained at a stimulus intensity of 30 V. Stimulus duration was constant at 0.2 ms. **C.** Representative field potentials recorded from hippocampal slices from tTA-KCC2 mice raised with Dox (DOX ON, upper traces) or with Dox withdrawn from the diet for 6-10 days (DOX OFF, lower traces), under control conditions and in the presence of different concentrations of the GABA_A_ receptor agonist muscimol. **D.** Muscimol concentration response curves for population spike amplitude in hippocampal slices from KCC2 and tTA-KCC2 mice, raised with Dox in the diet (Dox ON) or that had Dox withdrawn from the diet for 6-10 days (Dox OFF). Data is derived from 6 slices (from 4-5 mice) in each group. Curves represent the Hill equation fit to the mean data (r^2^ > 0.9).

### Increased KCC2 reduces seizure susceptibility in vitro

The above results suggest no differences in neuronal excitability or GABA receptor responses in the resting hippocampal slice. We next examined if upregulation of KCC2 impacted on neuronal excitability when neural circuit activity was increased by simulated seizure activity. Two different protocols were used, the first involved repeatedly applying a tetanus (100Hz, 1s) to the afferent pathways which could induce a spontaneous afterdischarge (AD) recorded in the CA1 pyramidal layer (Higashima *et al*., 2000). Tetani were applied every 10 minutes, and after the 3^rd^ or 4^th^ tetanus the AD pattern stabilized to a brief 5-20sec period of ADs consisting of typically 5-10 small (≈ 1 mV) voltage deflections (Fig. 5A). Such ADs were generally absent in slices from mice overexpressing KCC2 (Fig. 5A). The incidence of ADs was similar in the slices from the three control groups (KCC2 mice dox on 6/7 slices, KCC2 mice Dox off 6/7 slices, tTA-KCC2 mice Dox on 7/7 slices, 5 mice per group) and were of a similar number when present (Fig 5B). In contrast, ADs were significantly less common, only seen in one from 7 slices from five tTA-KCC2 Dox Off mice, giving rise to a significantly reduced mean AD frequency (Fig 5B). We also examined altered neuronal excitability in hippocampal slices perfused with Mg-free aCSF with elevated K^+^ (Mody *et al*., 1987). Perfusion of this Mg-free aCSF induced spontaneous spike discharges recorded at the extracellular electrode placed in the CA1 pyramidal cell layer (Fig 5C). In the three control mice cohorts, these spontaneous spikes were observed within about 5-10 mins after switching to the zero-Mg^2+^ solution, and stabilized at an approximate frequency of 0.2 to 0.5Hz (Fig. 5D). The latency to the appearance of spontaneous spikes was significantly longer in slices from KCC2 upregulated tTA-KCC2 Dox off mice, and the frequency of spontaneous spikes was significantly decreased in these slices (Figure 5D). In all slices, the spontaneous spikes rapidly disappeared once the perfusate returned to the standard, Mg^2+^-containing aCSF. Together these results clearly show that the development of hyperexcitability in hippocampal slices is significantly reduced by KCC2 upregulation.

**Figure 5.**
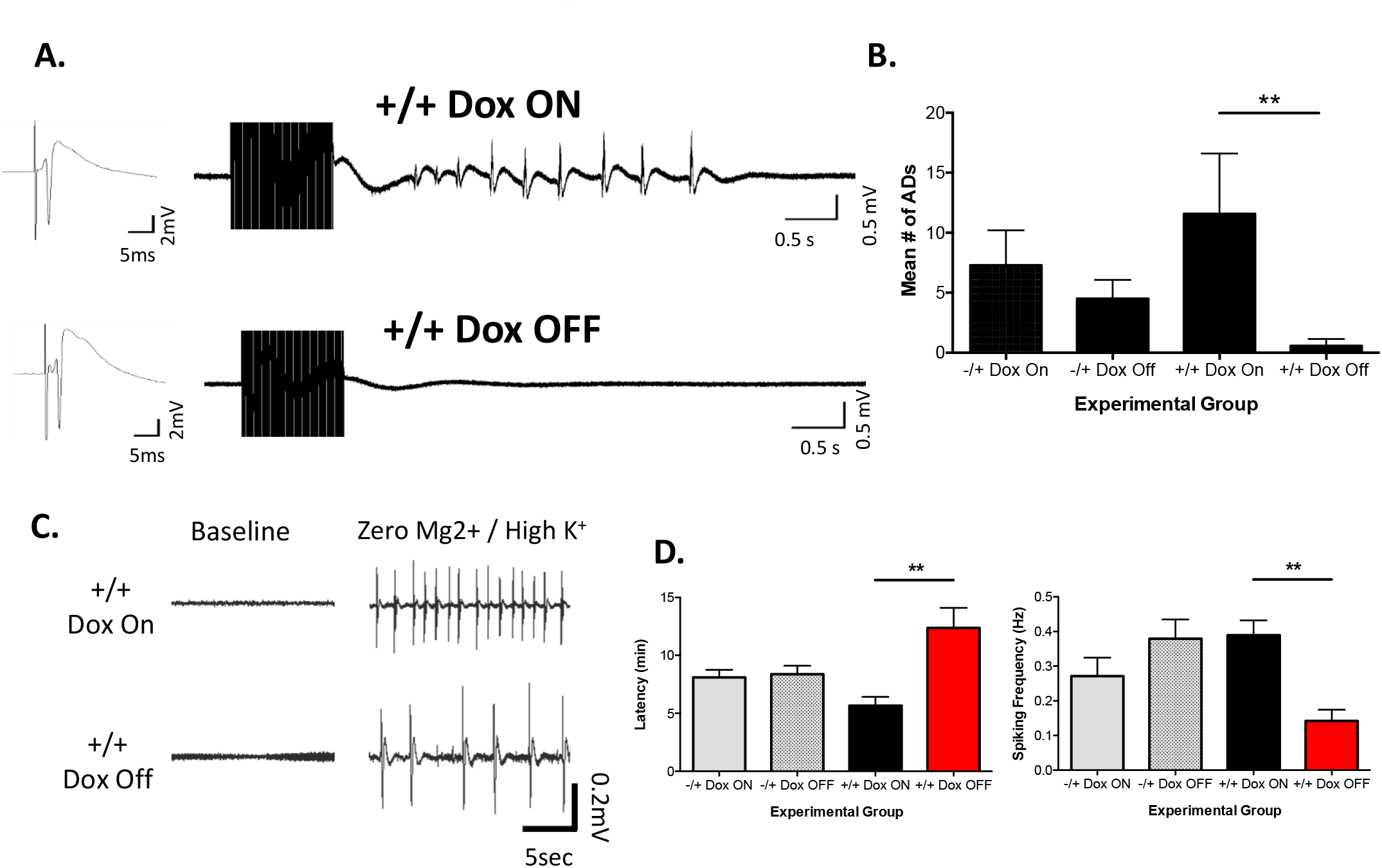
*Overexpression of KCC2 confers resistance to seizure activity in vitro.* **A.** Representative field potential responses recorded in CA1 cell layer in response to Schaeffer collateral stimulation. Left panels show a single stimulus evoked response. Right hand panels show the response to a 100Hz, 1sec tetanus. In slices from tTA-KCC2 mice raised with Dox (Dox ON) the tetanus induced spontaneous afterdischarges (ADs), which were absent in slices from Dox-off tTA-KCC2 mice. **B.** Plot of the mean number of ADs induced by tetanic stimulation, in slices derived from the mice groups as indicated (n= 6-7 slices from 5 mice in each group). The number of ADs in each slice was quantified by taking the average from the 4^th^ to 6^th^ tetanus (applied every 10 mins), by which time the response to the tetanus had stabilized. Overexpression of KCC2 resulted in a significant decrease in the mean number of ADs (** = p < 0.01; ANOVA and post-hoc Bonferroni’s test.) **C.** Representative field potentials recorded from CA1 pyramidal cell layer in control conditions (Baseline), and when Mg^2^+ was removed from the perfusate (Zero Mg^2+^ / High K^+^. Note the appearance of spontaneous discharges (spikes) upon perfusion with Mg^2+^-free solution. **D.** Quantification of the latency for appearance of spontaneous spikes in zero-Mg^2+^ perfusate (left) and the frequency of these spontaneous spikes (right). Data is derived from 6 slices (from 4-5 mice) in each experimental group as indicated. Overexpression of KCC2 resulted in a significant increase in spontaneous spike latency, and a significant decrease in the frequency of spontaneous discharges (** = p < 0.01); ANOVA and post-hoc Bonferroni’s test.

### Increased KCC2 prevents progression into Kainic Acid-induced Status Epilepticus

Kainic acid activates glutamate AMPA/kainate receptors and induces convulsions in rodents that can progress into a full status epilepticus (SE) (Ben-Ari & Cossart, 2000). To induce a more robust progression into SE without excessive mortality, and to also examine the threshold for SE more closely, we utilized a ramp-up dosing protocol and scored behavioural seizures using a modified Racine scale (McKhann *et al*., 2003; Tse *et al*., 2014). Mice were injected with a low dose of kainic acid (5mg/kg) every 30 mins until SE was observed, defined as a stage 4 seizure followed by continuous seizure behaviour or death. Figures 6A, 6B shows the patterns of seizure phenotypes in response to repetitive kainic acid doses for the tTA-KCC2 mice with Dox on and Dox off. At ≈ 15 mg/kg, the control mice cohorts (tTA-KCC2, Dox on; KCC2 Dox on and Dox off) typically experienced convulsive seizures with loss of posture, and progressed into SE (Figure 6A, 6C) whereas mice with upregulated KCC2 (tTA-KCC2) were relatively resistant to progression into SE, even when the dose was escalated to 50 mg/kg in total (Figure 6B, 6C). Indeed only 1 of five tTA-KCC2 Dox off mice experienced SE (at 50 mg/kg) whereas all 15 mice from the control cohorts experienced SE at total doses of 25 mg/kg or lower. This important result was replicated using the transgenic mice in Japan, prior to importation to Australia. In these experiments, 9 out of 10 control mice (either tTA-KCC2 Dox on or WT siblings) progressed into SE (at total doses up to 20-40 mg/kg) while none out of five tTA-KCC2 Dox off mice experienced SE at total maximal doses up to 30-50 mg/kg. In conclusion, the results clearly indicate that upregulation of KCC2 imparts a marked resistance to the progression into kainic acid induced status epilepticus.

**Figure 6.**
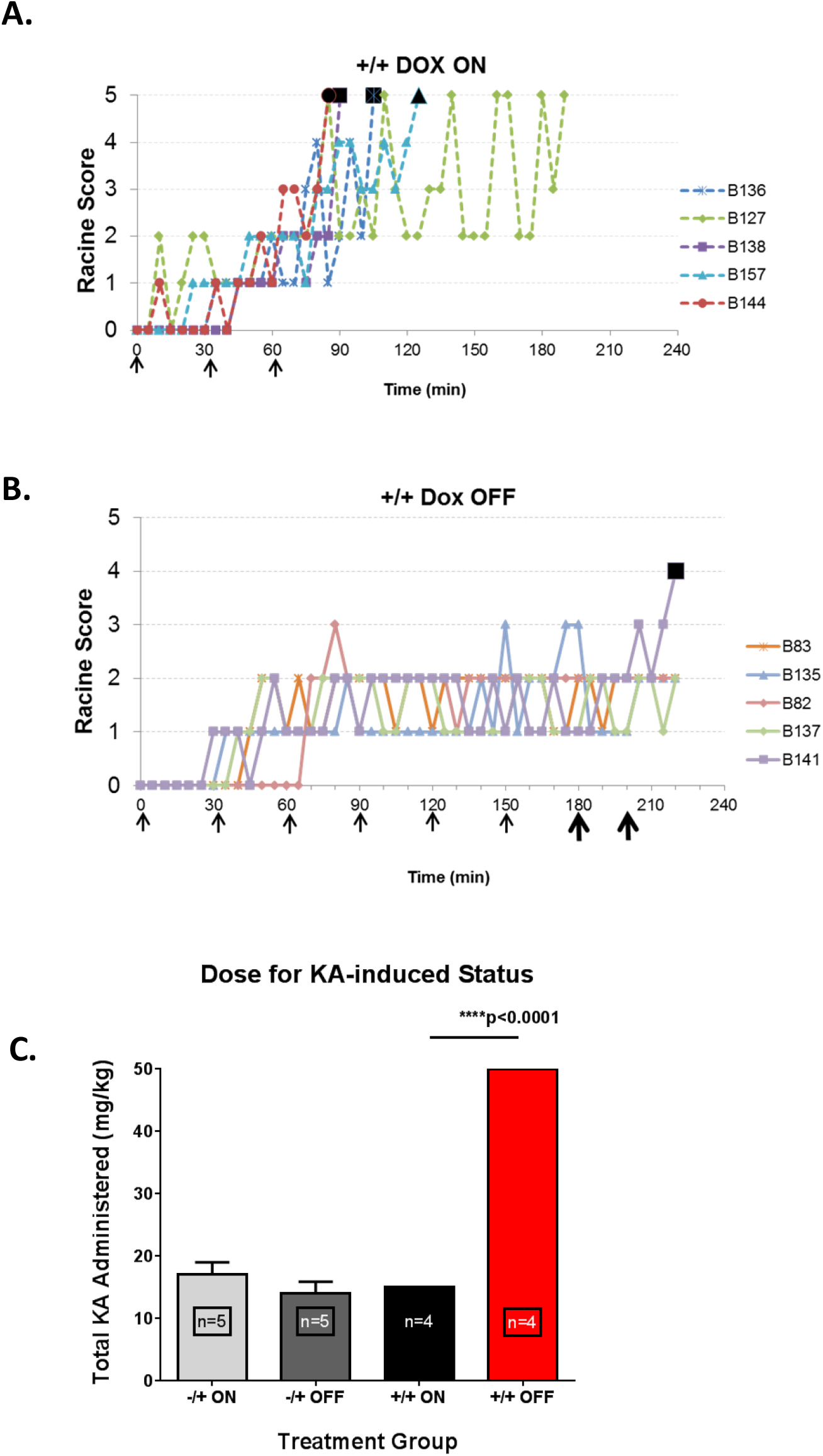
*KCC2 upregulation mitigates against the progression into Kainic Acid induced Status Epilepticus.* **A, B.** Representative plots of the timecourse of seizure phenotypes for each of five tTA-KCC2 (+/+) mouse raised on Dox (Dox on, **A**), and for each of five tTA-KCC2 (+/+) mice in which Dox was withdrawn from the diet for at least 6 days (Dox off, **B**). The time of KA injections (5mg/kg, ip) are indicated by arrow heads (bold arrowhead indicates 10 mg/kg, *ip*). Large bold symbol indicates seizure-induced mortality. **C.** Mean group data plotting the total dose of kainic acid required to elicit SE. The average dose required to elicit SE is indicated by the bar graphs (mean ± SEM), while data for from each individual mouse also shown. If no SE was observed after 50 mg total KA, this was assigned a data point of 50 mg. A significantly lower dose was required for SE in the Do-on tTA-KCC2 control mice as compared to the KCC2 overexpressing tTA-KCC2 Do off mice (*** p<0.0001, ANOVA and post-hoc Bonferroni’s).

### No effect of increased KCC2 on PTZ induced seizure responses

Finally, we investigated whether the seizure-resistance conferred by KCC2 upregulation in the escalating kainic acid challenge generalised across to other *in vivo* seizure models. The pentyltetrazole (PTZ) model is widely used to evaluate anticonvulsant drugs, with subcutaneous injections of this GABA_A_ receptor antagonist inducing a rapid progression through a well characterised series of behavioural seizure states. Mice from control (KCC2) and test (tTA-KCC2) genotypes, with and without Dox, were pre-administered either vehicle (30% ethanol in 0.9% saline, 0.1 ml, *ip*) or diazepam (3mg/kg in vehicle, ip) followed by a single dose of PTZ (85 mg/kg, *sc*). Seizures were scored blindly from video recordings based on a modified Racine scale of 6 stages (Luttjohann *et al*., 2009). In the vehicle cohort, all mice groups rapidly progressed through to higher grade convulsive and tonic seizures within 10-20 mins after injection. There was no significant effect of KCC2 upregulation on the response to PTZ. The latency to the first twitch seizure, and the total cumulative seizure score over the 30 mins post-PTZ is shown in Supplemental Figure 2A. Pretreatment with diazepam reduced the severity of the subsequent response to PTZ, as expected, but again there was no significant differences amongst the mice cohorts in the extent to which diazepam prolonged the seizure latency or reduced the total seizure score. In short, we could not detect any effect of KCC2 overexpression on the response to PTZ, nor on the effect of diazepam pre-treatment on the PTZ induced seizure response.

## Discussion

KCC2 plays a key role in neuronal Cl^-^ homeostasis and the efficacy of neuronal inhibition mediated by activation of GABA_A_ and glycine receptors in brain and spinal cord (Doyon *et al*., 2016). A downregulation of KCC2 function and expression occurs in a range of brain traumas and disease (Kahle *et al*., 2008; Kaila *et al*., 2014; Wu *et al*., 2016), including epilepsy (Cohen *et al*., 2002), chronic allodynia and spasticity from peripheral nerve injury (Coull *et al*., 2003; Boulenguez *et al*., 2010; Eto *et al*., 2012; Liabeuf *et al*., 2017), axotomy (Nabekura *et al*., 2002) and ischemia (Papp *et al*., 2008). Furthermore, alterations in Cl^-^homeostasis have been associated with cognitive deficits in autism spectrum disorders, Downs Syndrome, schizophrenia and age-related dementia (Deidda *et al*., 2015; Merner *et al*., 2015; Ferando *et al*., 2016; Tang *et al*., 2016; Ben-Ari, 2017). Consequently, intense interest has developed in manipulating KCC2 activity to potentially treat a range of different brain disorders. We have therefore established a conditional transgenic mouse to allow the selective overexpression of KCC2, and to facilitate the study of how increased KCC2 effects neuronal excitability and brain function *in vivo.*

Withdrawal of doxycycline from the diet lead to a rapid and reversible increase in KCC2. Within two days of withdrawing doxycycline, KCC2 mRNA and protein levels were increased. Protein levels increased by 2-3-fold by ≈ 7 days after Dox withdrawal. When neurons were challenged with an ionic load (NH_4_Cl), enhanced NH_4_^+^ membrane transport was seen in slices from transgenic mice, so at least some of this overexpressed KCC2 is functional. The increase in functional transport in this fluorescent assay was modest, although these fluorescent responses were averaged across multiple CA1 neurons within a slice, and not every cell would be expected to over express KCC2. In a series of transgenic mice lines, in which the tTA-tetO system was coupled to drive GFP fluorescence, between 30-70% of CA1 neurons failed to express GFP (Wu *et al*., 2016). The cellular extent to which KCC2 is overexpressed in our current tTA-KCC2 mouse after Dox withdrawal remains to be quantified.

The increase in functional KCC2 was sufficient to reduce the progression into seizures, both *in vitro* and *in vivo.* A striking effect was seen in two *in vitro* models: tetanic-induced afterdischarges were virtually absent while spontaneous spikes following perfusion with 0 Mg^2^+ hyperexcitable solution took longer to be induced and were of reduced frequency. This altered seizure threshold was seen in the absence of any major effects on basal population spikes or on GABA_A_ receptor responses in non-seizure or “basal” conditions. We propose that under such basal or resting conditions, neurons have sufficient functional KCC2 (and other transporters) to regulate intracellular Cl^-^ effectively, and additional KCC2 transporters have minimal impact on resting membrane potentials or excitability. Under resting or approximate equilibrium conditions, then the [Cl^-^]i set point should be established by the relative permeability for Cl^-^ and the driving forces for the ions that are actively transported across the membrane (e.g., E_K^+^_, E_CI^-^_ and E_Na^+^_ for KCC2 and NKCC1), with possible contributions from impermeant anions (Glykys *et al*., 2017). Increasing the relative number of KCC2 transporters present doesn’t appear to alter this set point. However, when the neural circuits become more active, such as following tetanic stimulation or when made hyperexcitable by 0 Mg^2+^, then we propose that endogenous KCC2 cannot maintain [Cl^-^] homeostasis, [Cl^-^]i becomes elevated, GABA inhibition is consequently weakened, and this contributes to spontaneous hippocampal discharges. When KCC2 is overexpressed, [Cl^-^]i and hence GABA inhibition is maintained during circuit hyperactivity, and seizures are reduced. A similar mechanism is proposed to account for the *in vivo* results. Under basal or non-seizure conditions, there is little effect of further increasing KCC2 transport on neuronal inhibition, and no sedation nor clear effects on locomotion or exploratory behaviour were seen in mice overexpressing KCC2. However, once the mice are challenged with by escalating kainic acid doses - then neural circuits become hyperactivated, [Cl^-^]i builds up, and inhibition becomes less efficacious. A build-up of [Cl^-^] in pyramidal neurons occurs during seizures and may trigger loss of functional inhibition and ictal events (Lillis *et al*., 2012; Glykys *et al*., 2017; Sulis Sato *et al*., 2017), as our data also supports. We propose that enhancing KCC2 enables better control of [Cl^-^]i and maintenance of neuronal inhibition. The initial and more modest seizures don’t seem to be prevented by KCC2 upregulation, but the progression of seizures into a sustained, self-propagating status epilepticus was prevented by increasing KCC2. Consistently, optogenetic *in vivo* activation of interneurons in the subiculum during the early stages of epileptogenesis can inhibit seizures, but their activation after GABA transmission has shifted to a depolarizing phenotype becomes less effective and can even exacerbate seizures (Wang *et al*., 2017). Hence enhancing Cl^-^ extrusion capacity may not prevent seizures, but may reduce the progression to a self-sustaining state associated with depolarizing GABA and loss of inhibition. This supports previous conclusions derived from the characterisation of the KCC2 S940A transgenic mouse, which mimics the specific downregulation of KCC2 during hyperactivity (Silayeva *et al*., 2015). Both *in vitro* and *in vivo*, seizure latency was unaffected, but the seizure severity was exacerbated with the greater loss of KCC2 membrane function. Similarly, blocking NKCC1 mediated Cl^-^ influx with bumetanide doesn’t prevent seizures, but can reduce their duration and severity (Sivakumaran & Maguire, 2016). Our data support the idea that enhancing KCC2 expression and function can be protective against seizures, and particularly in curbing their progression and severity. However, the generalisation of this neuroprotective strategy may be questioned by the complete lack of any protection seen in the PTZ models. Although PTZ and kainic acid induce seizures via different cellular targets, agents potentiating GABA_A_ receptors are effective in reducing PTZ seizure severity (as we also reported for diazepam). It may be that the rapid escalation of seizures in response to PTZ overwhelmed the capacity of the overexpressed KCC2 to maintain Cl^-^ homeostasis and robust inhibition. Regardless, the result highlights that enhancing KCC2 may not be a panacea to reduce all forms of seizures or epilepsy.

In conclusion, we present a conditional transgenic mouse in which functional KCC2 overexpression can be rapidly induced by dietary manipulation. Increasing KCC2 had little detectable effects on excitability or behaviour under non-pathological, resting conditions, strengthening its appeal as a potential therapeutic target and consistent with studies using pharmacological enhancement of KCC2 (Gagnon *et al*., 2013; Liabeuf *et al*., 2017). Overexpressing KCC2 was able to reduce the severity of seizures, indicating that neuronal inhibition may be selectively enhanced during hyperactive states by KCC2 upregulation. Our results suggest that the loss of KCC2 that can accompany seizures and epilepsy may be detrimental to restoring neural circuit homeostasis. Our mouse should facilitate investigations into the consequences of modulating KCC2 in a broad range of pathophysiological situations, helping guide potential therapeutic applications of targeting Cl^-^homeostasis to treat brain disorders.

## Methods

Experiments were approved by the UNSW Australia Animal Care and Ethics Committee and by the National Institutes of Natural Sciences of Japan, and adhered to the Guidelines for Animal Care and Welfare of the National Health and Medical Research Council of Australia.

### Generation and Maintenance of the Transgenic mouse colony

We first generated a KCC2 STOP-tetracycline operator (tetO) knock-in mouse (Tanaka *et al*., 2010). Homologous recombination was used to insert the KCC2 gene (from mouse BAC RP23-6103, Invitrogen) into a loxP-Neo-STOP-tetO plasmid (a kind gift from Dr kenji Tanaka, (Tanaka *et al*., 2010) which was electroporated into 129Sv/EvTac ES cells. Targeted ES cells were injected into C57Bl6J mice to generate a KCC2 STOP-tetO knock-in mouse. These mice were then crossed with ROSA-FLiP mice to excise the neoSTOP plasmid cassette to generate a KCC2-tetO mouse, with tetO inserted upstream of the KCC2 translation initiation site. KCC2-tetO mice were crossed with a calcium calmodulin-dependent kinase IIα (CAMKIIα)-tetracycline transactivator (tTA) mouse, which expresses tTA in forebrain neurons ((Mayford *et al*., 1996); a kind gift from Dr Kenji Tanaka). Mouse were genotyped from tail tips using separate PCR primers against the CAMKIIα-tTA and KCC2-tetO inserts. The forward and reverse primers for tTA were 5’-AGGCTTGAGATCTGGCCATAC-3’ and 5’-AAGGGCAAAAGTGAGTATGGTG-3’, respectively, and the KCC2-tetO forward and reverse primers used were: 5’-AGCAGAGCTCGTTTAGTGAACCGT-3’ and 5’-TGGAAACTCAAAGCGAGGAACTGC-3’ respectively.

Transgenic mice were maintained in both Japan and Australia, housed in a 12-12 light cycle with food and water ad libitum. Experiments were undertaken in Australia and/or Japan as indicated. Mice were raised and fed on doxycycline-laced chow (Australia: 600 mg/kg; Gordons stock suppliers, Narrabri NSW; Japan: 100mg/kg; Oriental Yeast Co., Ltd, Tokyo) and Dox off mice were separated and fed with the same Dox-free chow for 6-10 days prior to experiments, or as indicated.

### Molecular and biochemical validation

In situ hybridization (Japan; Fig. 1) was used to confirm overexpression of the KCC2 transcript. Adult (2-3 month) CamKIIα-tTA or tTA-KCC2 mice, either Dox on or Dox off were cardiac perfused with PBS followed by 4% PFA in PBS and brains excised and postfixed overnight, before cryoprotection in 20% sucrose and embedding in OCT compound. The methods are described in detail in Ma *et al*., (2006). Sagittal cryosections (30uM) were washed and prepared as described (Ma *et al*., 2006) and hybridized with digoxigenin-labelled KCC2 cRNA probes (NCBI accession number NM_020333.2; corresponding to 3’-untranslated region 3981-5993 bp; coding region is 416-3763 bp). Colour substrates (NBT/BCIP; Roche) were used for colour development, and nuclear fast red (Vector lab, Burlingame, CA) was used for counter-staining.

To quantify KCC2 protein expression using Western blots, two approaches were used. In the first (Japan; Fig. 1; (Watanabe *et al*., 2009)), brains were excised from adult male and female mice (2-3 months) and sliced at 300 μM using a Vibratome (Leica) in cold-aCSF. Regions of interest (1 mm diameter) were micropunched from each hemisphere, grouped and homogenized in lysis buffer (pH 7.4) on ice, containing Tris HCl (50 mM), NaCl (150 mM), EDTA (5 mM), complete protease inhibitor (Roche, Indiannapolis), 1% (w/v) Triton X-100. Homogenate was centrifuged at 12,000 G for 10 mins at 4°C and supernatant was collected. Proteins were separated in 7.5% acrylamide gels transferred to Immobilon-P membranes (Millipore, Bedford, MA). Reactions were blocked in 1% bovine serum albumin and membranes incubated overnight with the KCC2 (1:1000, Upstate Biotechnology, New York, now Millipore; Merck) or β actin (1:10000, Sigma-Aldrich) primary antibodies at 4°C. Blots were then incubated with HRP-conjugated secondary antibody (GE Healthcare UK Ltd., Buckinghamshire, England) for 1 h at room temperature and labelled protein visualized using enhanced chemiluminescence (ECL, GE Healthcare). Optical densities were quantified using Image J (NIH, Bethseda).

In the second method (Australia; Supp Figure 1), brains from adult KCC2 (control samples) and tTA-KCC2 mice, at different times after Dox withdrawal as indicated, were rapidly excised on to ice, and the hippocampi, cortices and cerebellum were dissected out and frozen for later homogenisation in 400ul RIPA lysis buffer (Cell Signaling) in the presence of protease (Sigma-Aldrich) and phosphatase (Roche) inhibitors. The supernatant was collected after centrifugation at 13000 rpm for 20mins, 4°C. Protein concentrations were determined with Biorad BCA assay and samples were denatured at 70°C for 10minutes. 10ug RIPA extracted protein was separated using 4-12% Bis-Tris acrylamide gel, and transferred to PVDF using the iBlot dry transfer system for 9mins, blocked in 5% skim milk and probed with polyclonal rabbit anti-KCC2 antibody (1:1000; cat&# 07-432; Upstate Biotechnology; Millipore), mouse anti-human transferrin receptor (1:1000; 13-6800; Invitrogen) or a monoclonal specific βactin (1:35 000; A3854; Sigma Aldrich) primary antibody overnight at 4°C. Blots were incubated with either peroxidase labelled anti-rabbit igG (1:10 000, Cell Signaling) or peroxidase labelled anti-mouse IgG (1:10000, Cell Signaling). Immunoreactive proteins were visualised using enhanced chemiluminescence (Thermo Scientific). Optical density measurements were performed using NIH Image J software.

### Imaging KCC2 transport in vitro using BCECF-AM

Mice (6-18 weeks; Japan) were anaesthetized with isoflurane and subsequently pentobarbitone (≈100mg/kg) and cardiac perfused with a cold, low Na+ cutting solution containing (in mM: Sucrose 248; MgCl_2_ 1; CaCl_2_ 1; MgSO_4_ 1.25; NaH_2_PO_4_ 2; KCl 2; NaHCO_3_ 26; D-Glucose 10; 95% O2 and 5% CO2. Brains were rapidly excised and sliced at 300-350 uM in the coronal plane (Leica VT100S), and then incubated in the standard, recording aCSF containing (in mM; NaCl 150, KCl 5, CaCl_2_ 2, MgCl_2_ 2, NaHCO_3_ 25, bubbled with 95% O_2_/ 5% CO_2_, pH 7.4). After at least 1 hour, a single slice was transferred to a small loading chamber filled with aCSF and 50-100 μM of BCECF-AM (2′,7′-bis-(2-carboxyethyl)-5-(and-6)-carboxyfluorescein) - acetoxymethyl ester; Invitrogen) along with some of the DMSO solvent (final volume 0.2-0.5% DMSO vol/vol) and supplemented with 0.1% (vol/vol) pluronic acid. Slices were loaded for 30-40 mins and then transferred to the recording chamber mounted on the stage of an Olympus (BX61WI) multi-photon microscope (objective: 20X 0.95 NA) and continuously perfused with aCSF at room temperature. BCECF fluorescence was excited using a Ti:Saphire mode-locked laser tuned to 790 nm, with emissions collected above 535 nm. A small region within the pyramidal cell layer, typically the CA1 subfield, with dye labelled neurons was identified and checked for fluorescent stability. Loading was variable and typically less than 10 fluorescent neurons were observed in a single field. A K^+^-free, NH_4_Cl (10 mM) containing aCSF was bath applied for 3-5 minutes with images continuously acquired (Fluoview ver 4.1a) at a frame rate of 1.4 to 2.5 Hz. Slices were then perfused with 0.5 mM furosemide for at least 8 minutes and the application of the NH_4_Cl solution was repeated in the continued presence of furosemide. In some slices a further 10 minute period of washout of furosemide was followed by a 3^rd^ application of NH_4_Cl. All solutions contained bumetanide (10 μM) to negate any contribution of NKCC1 to the fluorescent signal. Image files were imported into Image J for analysis. Between 2-8 individual neurons were marked as regions of interest (ROIs) and their fluorescent signal intensity (FI) was quantified and normalised to that observed just prior to the start of NH_4_Cl application. The relative fluorescence in each ROI was averaged to give a single trace for each slice. The peak relative FI during NH4Cl, and the minimum relative FI which occurred 2 minutes following NH_4_Cl application, under control conditions and with furosemide, was quantified and compared across mice. For animals maintained on Dox diet, Dox was added to ACSF at 6ng/mL.

### In vivo behaviour: Open field and Elevated Plus Maze

Male mice (11-18 weeks; Japan) were singly caged and transferred into the behavioural test room where they were acclimatized for 1-1.5 hours. Behavioural testing was performed between 3pm and 8pm, with open field performed on day 1 and elevated plus maze performed on day 2. Both control (wild-type and tTA mice) and tTA-KCC2 mice were raised with Dox chow, which was withdrawn in the Dox off cohort 6-8 days prior to behavioural testing. The open field chamber consisted of an opaque Perspex square with equal sides of 40 cm, with the centre area (24 x 24 cm) comprising 36% of the field area. A single mouse was placed into the same corner and activity was recorded over 10 minutes. The elevated plus maze comprised of two raised and bisecting 130 cm long laneways of 10 cm width with walls of 10 cm height. One of the laneways had clear Perspex walls in each arm, defined as the open arm. An enclosed central area extended 5 cm into each are, resulting in closed and open arms of 60 cm each. A single mouse was placed into the central area facing towards the same closed arm, and its activity was monitored for the subsequent 10 minutes. One mouse jumped off the open arm during the trial and data for this mouse was excluded from further analysis. Ambient temperature and lighting was 25-30°C and typically 100-150 lux, and apparatus was wiped clean with 70% ethanol or hyperchlorous solution between each trial. Movies were captured via a USB camera (Logicool HD720p, Logitech, Japan) mounted above the behavioural apparatus and analysed later by an observer blind to the genotype using Anymaze software (Woods Dale, Illinois). The centre of the mouse was used to determine the mouse’s position, as this was identified in a subset of data as most consistently matching the manually scored data. Contrast was adjusted to minimize false detection events and all analysis was visually inspected to confirm valid scoring events.

### In vivo behaviour: Pentyltetrazole and Kainic Acid assays

For the pentylenetetrazole (PTZ) induced seizure assay (Australia), mice of both sexes (8-14 weeks, gender balanced) were transferred to the recording room and acclimatized to the recording chamber for 30 minutes prior to starting experiments. The experimental chamber consisted of a clear Perspex box (15 cm width x 25 cm long x 15 cm high) into which were mounted a series of mirrors so that detailed observations of mouse facial features and behaviour could be recorded with an HD video camera mounted to the side of the chamber. 15 mins prior to PTZ injection mice were administered either vehicle (30% ethanol in 0.9% saline, 0.1 ml, sc) or diazepam (3 mg/kg, 0.75 mg/mL of vehicle, sc). PTZ (85 mg/kg in saline) was administered as a single *sc* bolus, and behaviour was monitored for the appearance of a stage 5 seizure or for up to 60 mins, after which surviving mice were euthanized and tissues extracted. Video recordings were coded and scored later in a blind fashion according to a modified 6 stage Racine scale for PTZ (Luttjohann *et al*., 2009), where stage 3 and higher represented convulsive or tonic-clonic seizures. The latency to Stages 3 and 5 were quantified, and the dominant Stage each 5 minutes was tabulated to give a cumulative seizure score.

For the Kainic Acid (KA) induced seizure assay (Australia), male and female mice (8-14 weeks, gender balanced) were singly caged in the experimental room and left to acclimatize for 60 mins. Mouse behaviours were monitored by a camera mounted in front and above their cage and the experimenter was blinded to the genotype. Mice were injected with KA (Sigma, St Louis) using a dose escalation paradigm in which a modest does of 5mg/kg was administered *ip* every 30-40 mins until a stage 4 or 5 seizure was observed. Seizure scoring used again a modified Racine scale characterised as follows: Stage 1, immobility; Stage 2 Straub tail, piloerection; Stage 3 brief convulsive seizure (twitch) and/or forelimb cycling; Stage 4 additional loss of posture, Stage 5 jumping, bouncing seizure. Stage 4-5 seizures were typically followed continued episodes of stage 3-5 seizures that could be interspersed with stage 1-2 seizures and that lasted at least 60 mins, although mice sometimes died during this SE and hence SE was defined here by the presence of at least two Stage 4 seizures. Dosing continued until SE was reached or until a total of 50 mg/kg had been administered. These experiments were also repeated with the Japanese colony, with a similar dosing escalation schedule and scoring system, except the experimenter was not blind to the genotype. Also, test and sibling controls were tested concurrently under the same conditions.

### Hippocampal slice electrophysiology

Mice (Australia) of either sex (8-16 weeks) were anaesthetized with pentobarbitone (50 mg/kg, ip) and cardiac perfused with an ice cold, sucrose aCSF solution containing (in mM): Sucrose, 230; MgCl_2_, 1; CaCl_2_, 0.5; KH_2_PO_4_, 1; KCl, 2; NaHCO_3_, 26; D-Glucose, 10; 95% O_2_ and 5% CO_2_. Brains were rapidly excised and sliced at 400 μM in the sagittal plane (Leica VT1200) in the cold cutting solution before being transferred to a slice incubation solution filled with aCSF (in mM; NaCl, 125; KCl, 2.5; NH_2_PO_4_, 1.25; CaCl_2_, 2; MgSO_4_, 1; NaHCO_3_, 25; bubbled with 95% O_2_/ 5% CO_2_, pH 7.4). *In vitro* seizure experiments were conducted at 32 °C, while muscimol concentration responses were done at 24 °C. The slice incubation chamber was set at the same temperature as the recording chamber. A single slice was transferred to the recording chamber and extracellular field potential responses were recorded from the CA1 region of the stratum pyramidale in response to stimulation of Schaeffer collateral afferent fibres. The recording electrode was a glass pipette filled with aCSF and the bipolar stimulating electrode was constructed of fine (25 μm) Ag wires insulated up to the tip with Teflon connected to a constant current digital stimulus isolation unit (DS2A-Mkii, Digitime**r**). Acquisition was via an Axoclamp 900A (Molecular Devices), PowerLab (4/25T) and LabChart™ software (v7, ADInstruments), acquired at 5kHz following low-pass and high pass filtering at 0.1Hz and 2kHz, respectively (Powerlab, AD Instruments).

Once stable recordings were obtained from a slice, a series of single stimuli were applied to determine the stimulus intensity that produced a maximal and half maximal population spike (PS). Baseline responses were obtained at 0.1 Hz and only slices in which a clearly identified PS with a maximal amplitude of > 1 mV were used for experiments. For concentration response experiments, stimuli at an intensity to produce an approximate 50% maximal response were applied continually at 0.1 Hz and different concentrations of muscimol were sequentially bath applied for at least 30 mins at each concentration. A set of control recordings were interspersed between every 2-3 muscimol concentrations, allowing at least 40 minutes for drug washout. Between 4-6 concentrations were obtained for each slice, and control responses flanking each concentration were averaged and used for normalisation of the data. PS amplitude was measured by interpolating the response amplitude between before and after the PS, and measuring from this extrapolated baseline to the response peak. Normalised data from each experiment was grouped and fit to the Hill equation using Prism™ (Graphpad, CA) to obtain the maximal inhibition, IC50 and Hill slope. Fits to the sets of data from different mice groups were done concurrently and constrained to have the same slopes. For the input-output curves, at least 3 evoked responses at each stimulus intensity were averaged to derive the PS amplitude value at that intensity. Responses were then normalised to the maximal PS amplitude for that slice, allowing for averaging of multiple slices within in each experimental group.

To evaluate neuronal excitability in hippocampal slices, two seizure models were used. In the first (Higashima *et al*., 2000), control PS responses were initially evaluated to ensure stability and then a 10 second tetanic stimulus train at 100 Hz was applied to the Schaeffer Collaterals at a stimulus intensity that was just supramaximal. Stimulus trains were repeated every 10 minutes for 1 hour and the presence and number of afterdischarges were manually quantified. In the second protocol (Mody *et al*., 1987), control PS responses were again initially evaluated for amplitude and stability and then the bath was perfused with a zero Mg^2+^ solution consisting of (in mM): NaCl, 125; KCl, 5; NH_2_PO_4_, 1.25; CaCl_2_, 2; Glucose,10; NaHCO_3_, 26, which produced spontaneous field potential transients (spikes) at the CA1 recording electrode. The latency to the appearance of these spontaneous spikes, and the frequency of these spikes was manually quantified. Spike frequency was determined from the time of 1st spontaneous spike till the end of the perfusion period

For animals maintained on Dox diet, Dox was added to ACSF at 6ng/mL.

### Data Analysis and Statistics

All data are expressed as mean ± SEM and were analysed using Prism™ (Graphpad, CA). Multiple genotypes and diet groups were compared using an ANOVA and with three post hoc comparisons preselected prior to analysis to maximize statistical power: i) Effect of Doxycycline Diet Alone: KCC2 Dox On vs. KCC2 Dox Off mice; ii) Effect of Genotype Alone: KCC2 Dox On vs. tTA-KCC2 Dox On mice; and iii) Effect of KCC2 Upregulation: tTA-KCC2 Dox On vs. tTA-KCC2 Dox Off mice.

---

**Supplemental Figure 1.**
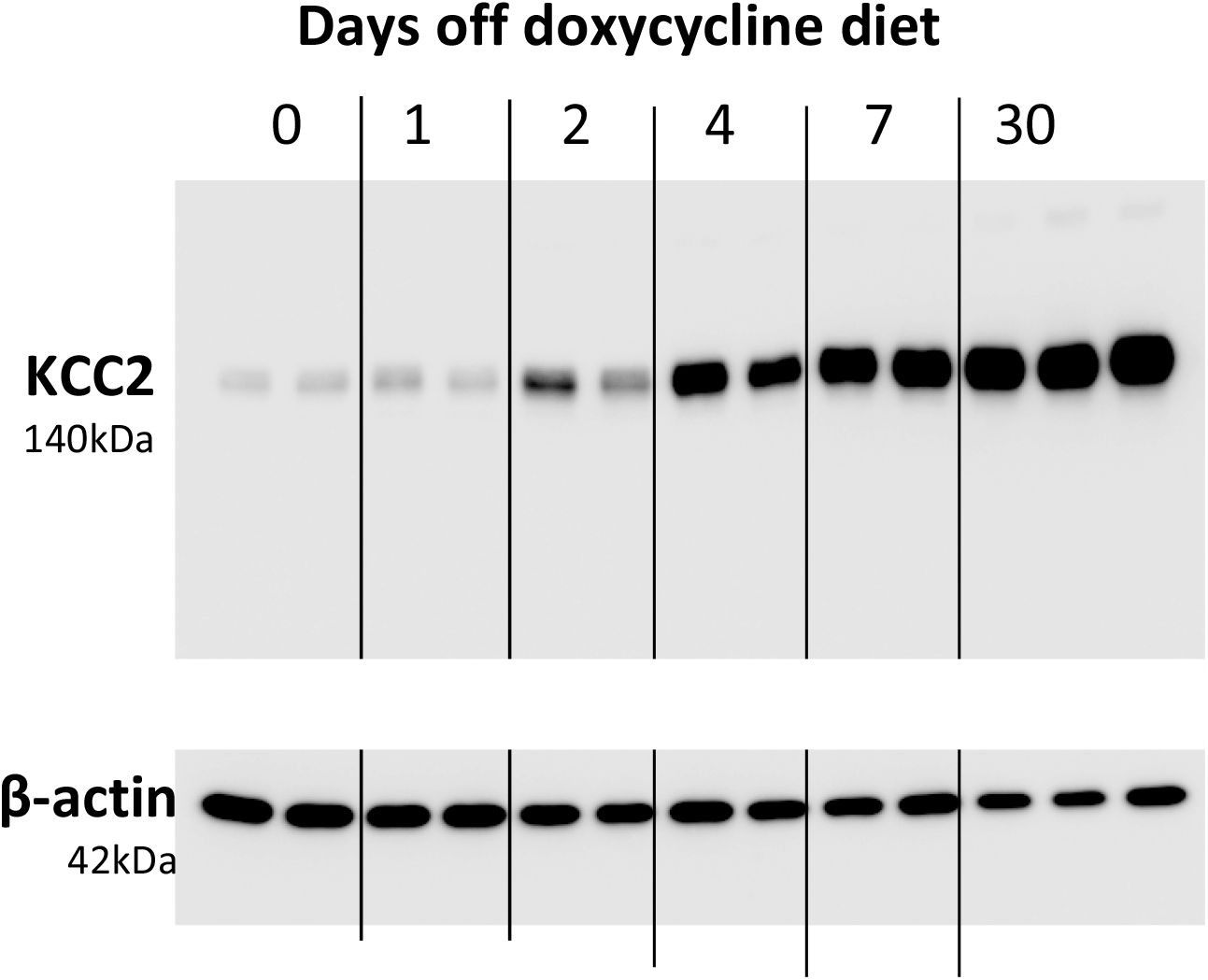
*Time-course of Dox withdrawal induced KCC2 overexpression.* Upper panel shows sample western blots for KCC2 in hippocampal tissues from 2 different tTA-KCC2 mice after withdrawing Dox from their diets for 0-30 days as indicated (0 days = Dox on). Lower panel shows immunoblot for beta-actin from same tissue samples.

**Supplemental Figure 2.**
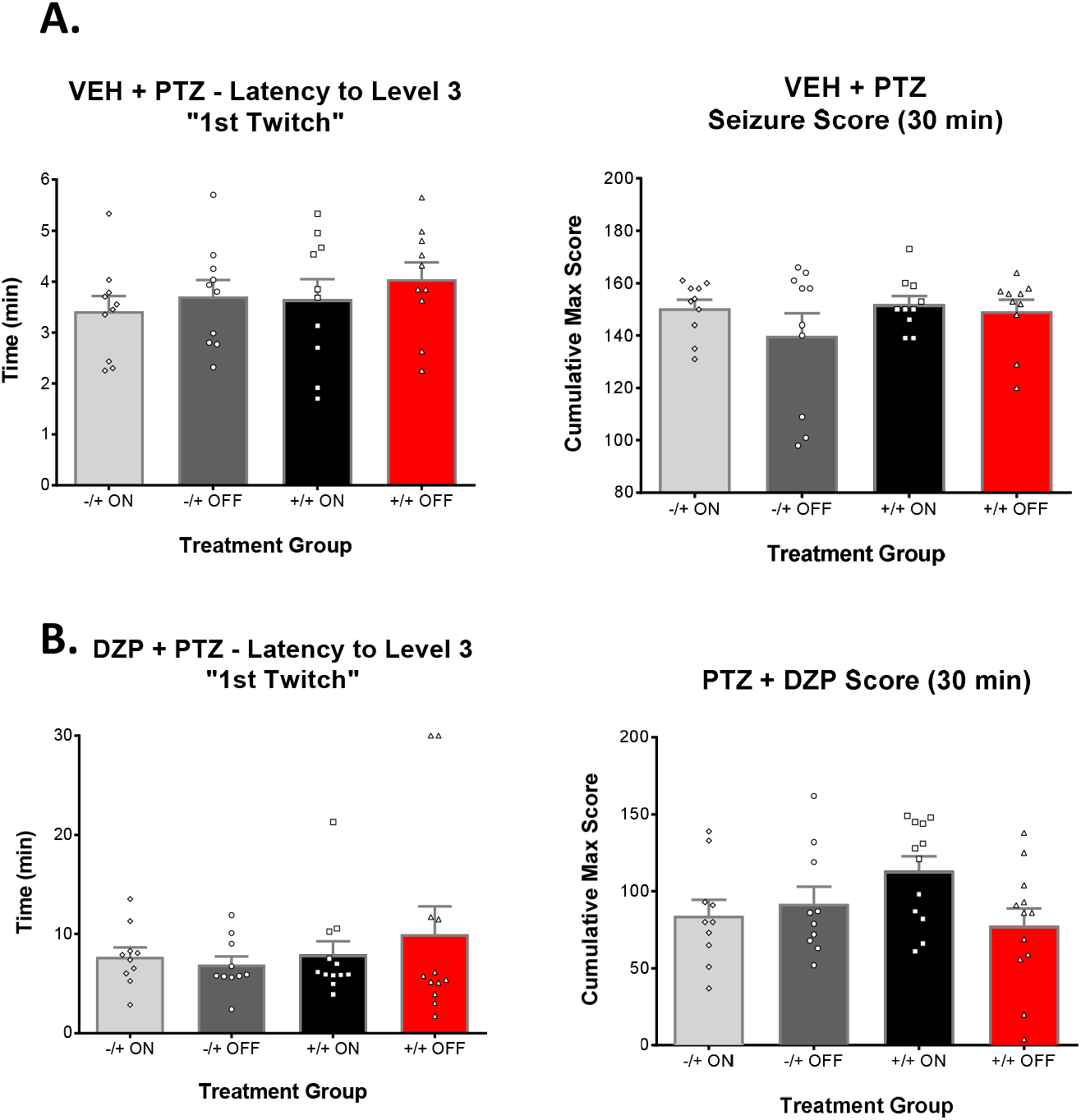
*Pentyltetrazole induced seizure phenotypes are unaffected by KCC2 overexpression* **A.** Mean group data showing the latency to the 1^st^ appearance of a generalized clonic seizure (“twitch”) in response to PTZ (85 mg, kg, sc) after saline (left) or diazepam (right) pretreatment. **B.** Mean group data showing the cumulative seizure score during the 30 (left) or 60 (right) mins following administration of PTZ (85 mg/kg, sc) with saline (left) or diazepam (right) pretreatment. In A and B, results from KCC2 and tTA-KCC2 mice, with and without Dox in the diet are shown (n= 10-13 in each group). Column graphs represent mean ± SEM, and data from each individual mouse also shown. There was no significant effect of KCC2 overexpression on any of these parameters (ANOVA and post-hoc Bonferroni comparison).

